# Evaluation of near infrared spectroscopy for sporozoite detection in mosquitoes infected with wild-strain parasites from asymptomatic gametocyte carriers in Kilifi Kenya

**DOI:** 10.1101/2020.07.25.220830

**Authors:** Marta F. Maia, Martin G. Wagah, Jonathan Karisa, Robert Mwakesi, Festus Mure, Martha Muturi, Juliana Wambua, Mainga Hamaluba, Floyd E. Dowell, Philip Bejon, Melissa C. Kapulu

## Abstract

**Background:** Screening for *Plasmodium* spp. sporozoite infection in mosquitoes is routinely done using ELISA (enzyme-linked immunosorbent assay). Near infrared spectroscopy (NIRS), a fast and non-destructive method, has recently been shown to distinguish, with 95% accuracy, between uninfected and sporozoite-infected mosquitoes using laboratory strains of *Plasmodium falciparum* (PfN54). The aim of this present study was to further investigate the reproducibility of NIRS to identify sporozoite infection in mosquitoes infected using field isolates of *P. falciparum* gametocytes from asymptomatic carriers.

**Methods:** Healthy individuals (aged 5 years and above) were screened for gametocytaemia by thick-smear microscopy in an area of moderate transmission along the Coast of Kenya between May and September 2018. Asymptomatic gametocyte carriers were recruited for mosquito feeding assays, direct membrane feeding (DMFA) and direct skin feeding (DFA), using insectary-reared *Anopheles gambiae* s.s (Kilifi strain). Mosquitoes were kept for 14-days post feeding after which they were scanned using NIRS and subsequently analysed for sporozoite infection using circumsporozoite-ELISA. Predictive models were explored using partial least square regressions (PLS).

**Results:** Two hundred and ninety-nine (299) individuals were screened for malaria parasites, 74 (24.8%) were found with circulating asexual parasites, and 16 (5.4%) with *P. falciparum* gametocyte stages. Fourteen (14) asymptomatic gametocyte carriers were recruited to the study for mosquito feeding assays. A total of 134 (7%, 134/1881) sporozoite-infected mosquitoes were obtained from 9 successful experiments. Three different training datasets composed of infected and uninfected mosquitoes were analysed. The PLS models were unable to distinguish between sporozoite-infected and uninfected mosquitoes. A predictive model could not be generated.

**Conclusions:** The results of this study were not consistent with previous published research on NIRS for detection of sporozoite infection in the same mosquito species and may reflect differences between laboratory and field conditions. The current findings indicate that methods for sporozoite detection should be tested on field isolates at an early stage in their development and are informative for future research seeking novel high-throughput methods for parasite detection in mosquitoes.

## INTRODUCTION

The WHO’s Global Technical Strategy for Malaria which seeks to reduce malaria incidence and related mortality by at least 90% and to eliminate the disease in a minimum of 35 countries by 2030 is off-track [1]. New control tools are needed and surveillance must be implemented as a core intervention to better inform malaria control programmes as well as the development of new control tools [2]. Human host parameters, such as asymptomatic community parasite prevalence, are widely used to reflect transmission intensity, but malaria transmission is most directly estimated using entomological parameters by measuring the proportion of sporozoite-infected Anopheline mosquitoes that attempt to bite humans in space and time. Screening for sporozoite infection in mosquitoes is routinely done using ELISA (enzyme-linked immunosorbent assay) [3], but it is a time-consuming and laborious process, hence our current reliance on indirect measures of transmission by monitoring infection in the host. New spectral methods such as NIRS (near infrared spectroscopy) and MALDI-TOF MS (matrix-assisted laser desorption/ionization time-of-flight) are currently being researched as potential alternatives [4] as they could provide high-throughput and lower cost per sample allowing surveillance and elimination programmes to screen large numbers of mosquitoes and thus better estimate the EIR (entomological inoculation rate), as will be required to assess progress towards malaria elimination [5].

NIRS has been applied to analyse various entomological traits in malaria vectors including species [6, 7], age [6, 7], and more recently *Plasmodium* spp. infection [4, 8]. Light in the visible and near infrared region of the spectrum (wavelength 400-2500 nanometers) is passed through a mosquito’s cephalothorax and an absorbance spectrum is collected instantly. The method is non-destructive and requires no consumables. As the external and internal biochemical composition of a mosquito changes so does its NIR absorbance spectrum. Consistent biochemical changes related to specific traits in mosquitoes can be used to develop predictive models. We have recently/previously shown that NIRS accurately predicted both oocyst and sporozoite infection in *Anopheles gambiae* s.s. mosquitoes that had been infected with laboratory-cultured *Plasmodium falciparum* (PfN54) gametocytes [4]. However, mosquitoes infected in the lab may be more alike and lack the natural variability present in mosquitoes infected in the wild and/or locally adapted laboratory strains, in addition the concentration of gametocytes ingested during a feed are considerably different between cultured and circulating gametocytes. For this reason, we designed a study to evaluate if NIRS would be able to distinguish between uninfected mosquitoes, and mosquitoes that had been infected after ingesting blood from asymptomatic malaria carriers.

## METHODS

### Study design

The study aimed at testing the hypothesis that NIRS could distinguish between uninfected and sporozoite-infected mosquitoes that had been infected with circulating *P. falciparum* strains. The target number of gametocyte carriers was calculated considering the number of mosquitoes required to generate and test a NIRS calibration. Based on other NIRS studies on insect traits [6], 100 sporozoite-infected and 100 uninfected mosquitoes were decided as minimum needed for testing (50-training dataset) and validating (50-test dataset) differences between the NIR spectra of the two categories, sporozoite-infected and uninfected. It was expected that the proportion of successful experiments, defined as at least one mosquito becoming infected after blood feeding would be around 40% [9], and that only 10-15% of engorged mosquitoes would become infected and survive 14 days post blood feeding [10]. Hence, it was concluded that a target number of 14 gametocyte carriers needed to be recruited for mosquito feeding assays in order to obtain sufficient number of infected mosquitoes to test and validate the NIRS method.

### Participant screening, recruitment and parasite detection

Asymptomatic gametocyte carriers, 5 years-old and above, were identified through house-to-house visits in various sub-locations of Kilifi South along the Coast of Kenya (Figure 1) between May and September 2018. Information on recent malaria cases from the surrounding health facilities were used to target homesteads where a member had been recently treated for malaria. Participants were allowed ample time to review the details of the study in the participant information sheet, upon which written informed consent was obtained for screening of asymptomatic individuals residing in the study area. Parental consent was obtained for all children (5-17 years old) and informed assent was additionally obtained from children between 12 and 17 years old.

**Figure 1.**
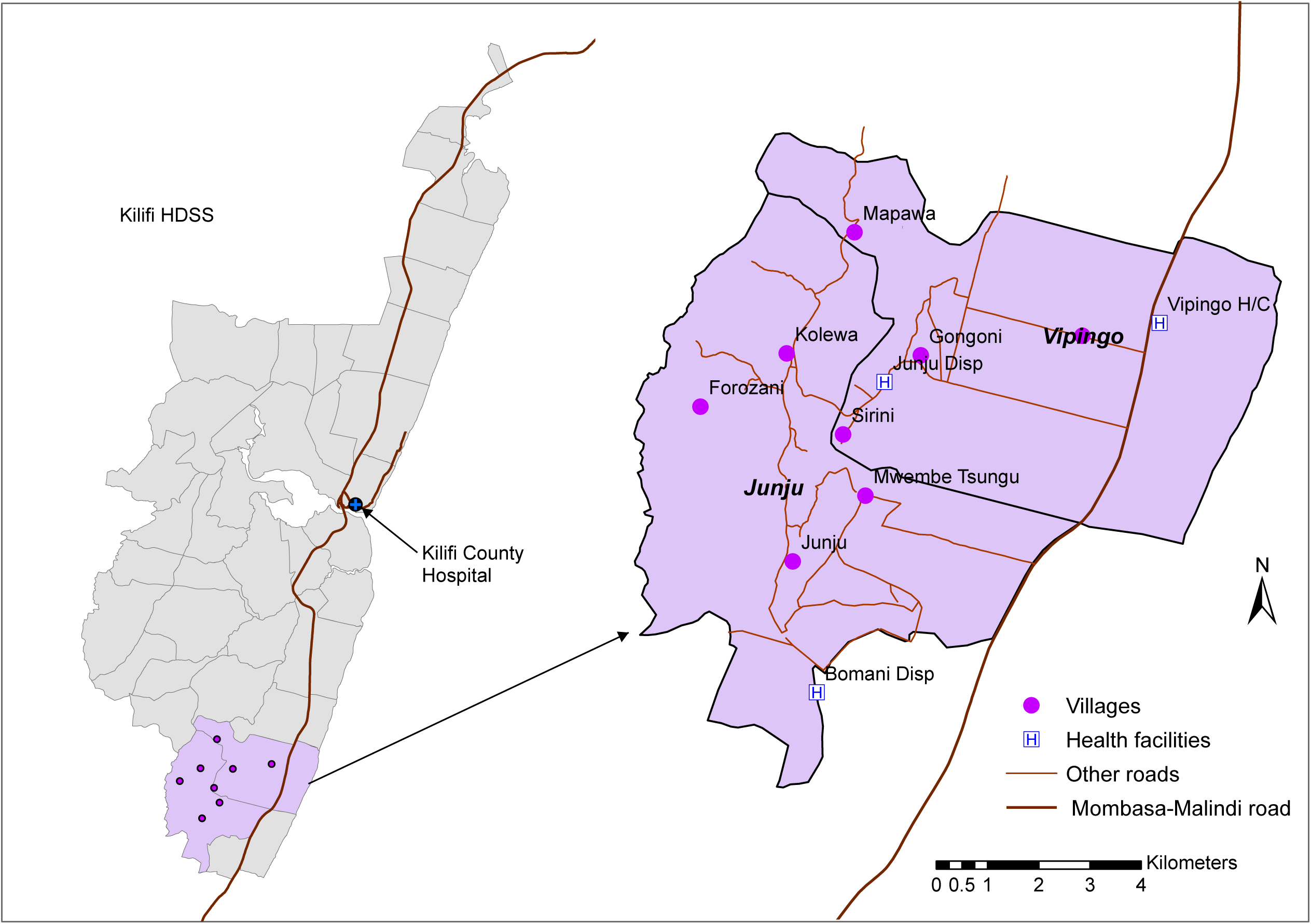
Map of the study area showing the location of the villages where participants were screened for malaria parasites between 28th May and 18^th^ September 2018.

Screening for circulating gametocytes was done by finger prick to prepare thick and thin blood smears in the field which were transported the same day to KEMRI Wellcome Trust Research Programme (KWTRP) in Kilifi town. Screening was done three days a week and no more than 20 slides were taken per day to KWTRP to be read by a trained microscopist. The thin blood films were fixed with 100% methanol and stained with 3% Giemsa stain for 45 minutes. Thick films were air-dried before staining. Thick films were first inspected, if there were more than 25 parasites per field on the thick film then the thin film was used for counting, otherwise the thick film was used. Parasite densities per mcl (microliter) of blood were calculated as the number of parasites per 200 white blood cells (WBCs) for thick films or per 500 red blood cells (RBCs) for thin films. The final parasite density was estimated assuming a WBC count of 8×10 ^9^ per Litre. The presence, parasite species, and densities of asexual and sexual parasites were recorded. A slide was considered negative for malaria infection after 100 fields of a thick film were read and no parasites were found. Parasite counts were determined by two independent microscopists and when necessary resolved by a third. Participants found with circulating asexual parasites were contacted by a field worker and referred to their local dispensary for treatment according to Kenyan national malaria treatment guidelines [11].

Individuals found with circulating gametocytes were also contacted by a field worker and asked to visit the KWTRP the following morning for participation in mosquito feeding assays. Once at KWTRP, participants were asked about recent medical history and medication, and axillary temperature was measured. A thick and thin film was repeated to determine gametocytaemia on the day of the feeding assays. Participants were excluded if they had taken anti-parasitic medication or antibiotics in the past two weeks, had a temperature over 37.5 degrees Celsius, felt unwell, or were found to no longer have circulating gametocytes.

### Mosquito colony maintenance

Mosquitoes from a colony of *An. gambiae* (Kilifi strain) were reared under standard insectary conditions (26±1°C, 80% humidity, 12 hr light: 12 hr dark cycle) at the KWTRP. Larvae were fed on Tetramin tropical flakes and pupae were transferred into cages for adult emergence. Adult mosquitoes were fed *ad libitum* on 10% glucose solution. Feeding assays were done using 2-5 days old mosquitoes previously starved of glucose and water for 8 h and which had never been given a blood meal (blood-naïve).

### Mosquito feeding assays

A trained phlebotomist withdrew 4ml of venous blood through venipuncture for direct membrane feeding assay (DMFA). Adult participants (18 years or older) were also asked if they would consent to directly feeding 50 mosquitoes on their forearm for 20 minutes through direct feeding assay (DFA). Participants could withdraw from either DMFA or DFA without need to provide justification.

DMFA was done at the KWTRP insectary using a water-jacketed membrane feeding system heated with a water bath kept between 38 degrees Celsius (C). The water-jacketed membrane feeding system was composed of a series of 4 mouth-blown glass feeders which were covered with Parafilm membrane on the bottom. The system was heated at least 30 minutes in advance of the DMFA to avoid temperature fluctuations in the glass feeders. All material used for the blood draw, including, syringe, butterfly and sample collection tube were kept at 38 degrees in a portable incubator and were only removed immediately before use. After blood was drawn, precautions were taken to not shake the sample tube and 900 microliters of whole blood were transferred to each glass feeder using a warm sterile pipette. Serum replacement was not done as it was the intention of the study to replicate as best as possible what happens in the natural environment with mosquitoes feeding on whole blood with host serum. A total of 4 cups were used per DMFA, each containing approximately 50 to 75 starved, blood-naïve, female *An. gambiae* s.s. mosquitoes amounting to between 200 and 300 mosquitoes per DMFA participant.

Participants older than 18, who volunteered to take part in the DFA, fed an additional 50 mosquitoes on their arm for 20 minutes. During DFA there was no risk of disease transmission to the participant as all mosquitoes were insectary-reared and blood-naïve.

After the mosquito feeding assays, the participants were referred to the Out-Patient Department of the Kilifi District Hospital (KOPD) where they were seen by a clinical officer and treated according to the Kenyan national malaria treatment guidelines [11]. If the participant had agreed to the DFA they were also prescribed an anti-histamine cream to relieve any pruritus caused by the mosquito bites.

### NIRS spectra collection

Blood-engorged mosquitoes were maintained in an access-controlled insectary for 14 days post blood-feeding to allow sporozoite development. After 14 days the mosquitoes were killed using chloroform vapors. Noticeably desiccated mosquitoes which had died earlier than day 14 were discarded. A near infrared (NIR) absorbance spectra was collected from each mosquito, without any further processing, using a Labspec 4i NIR spectrometer with an internal 18.6 W light source (ASD Inc, Longmont, CO) and ASD software RS3 and a 3.2 mm-diameter bifurcated fiber-optic probe which contained a single 600-micron collection fiber surrounded by six 600 micron illumination fibers. The probe was placed exactly 2.4 mm from a spectralon plate onto which the mosquitoes were placed for scanning. All mosquitoes were scanned on their cephalothorax and given a unique identifier.

### Sporozoite ELISA

After scanning, each mosquito carcass was stored individually at -80 °C until circumsporozoite enzyme linked immunosorbent assay (CSP-ELISA) was done for detection of *Plasmodium falciparum* circumsporozoite protein (CSP) in each individual mosquito. Standard ELISA protocol was followed [12] using *P. falciparum* Sporozoite ELISA Reagent Kit, MRA-890, containing lyophilized monoclonal antibody, were obtained through BEI Resources, National Institute of Allergy and Infectious Diseases (NIAID) and National Institutes of Health (NIH). Three negative controls and three positive controls were included per assay. Positive controls were provided with the kit, whilst the cephalothorax of male mosquitoes (*A. gambiae s*.*s*. from the same insectary) were used as negative controls.

### Data analysis

The results from the sporozoite ELISA were used to identify which individual mosquitoes had developed sporozoite infection. This information was specified to each respective NIR absorbance spectrum and added as a constituent to the predictive model. An equal number of spectra from uninfected and infected mosquitoes were randomly selected to explore differences in the NIR spectra between infected and uninfected mosquitoes. Leave-one-out cross validations (LOOCV) using partial least square (PLS) regressions in GRAMS Plus/IQ software (Thermo Galactic, Salem, NH) were used to analyze the two groups of spectra, and evaluate the predictive performance of the model.

LOOCV is a *k-fold* cross validation, with *k* equal to the total number *(n)* of spectra in a training dataset. That means that *n* separate times, the function approximator is trained on all the spectra except for one spectrum which is left-out. The algorithm then predicts the *left-out* spectra with the remaining spectra *(k-1)*, which, in combination, become the training sample. The out-of-sample predictive accuracy of the model is assessed by estimating the predicted residual sum of squares (PRESS) and the number of latent factors needed to explain the variation between spectra groups. R^2^ and predicted R^2^ are evaluated as measures of in-sample explanatory power. If a model performs well at *self-prediction*, a training dataset is then used to develop a calibration file to test the predictive ability of the model on an independent set of spectra (test dataset).

## RESULTS

### Parasitological survey

A total of 179 adults (18 years or above) and 120 children (5-17 years old) were screened from various sublocations of Kilifi South in Kilifi County (Figure 1). The highest number of parasite infected individuals were found in Junju, Kolewa and Mapawa sub-locations (Table 1). From the 299 screened participants, seventy-four (74, 24.8%) were found with circulating asexual parasites, and 16 (5.4%) with *P. falciparum* gametocyte stages (Figure 2). In eleven individuals (3.7%) both asexual and sexual parasites were found in circulation, of which ten were positive for *P. falciparum* and 1 for *Plasmodium ovale*.

**Table 1.** Number of participants screened and found to have circulating *Plasmodium spp* parasites in sublocations of Kilifi South, Kilifi County, Kenya.

| Village | Asexual parasites, n/N (%) | Gametocytes, n/N (%) | Total screened |
| --- | --- | --- | --- |
| Gongoni | 8/61 (13.1%) | 2/61 (3.3%) | 61 |
| Junju | 16/66 (24.2%) | 4/66 (6.1%) | 66 |
| Kolewa<sup>1</sup> | 21/58 (36.2%) | 3/58 (5.2%) | 58 |
| Mapawa | 24/88 (27.3%) | 5/88 (5.7%) | 88 |
| Mwembe Tsungu | 4/17 (23.5%) | 3/17 (17.6%) | 17 |
| Sirini | 0/7 (0%) | 0/7 (0%) | 7 |
| Forozani | 0/1 (0%) | 0/1 (0%) | 1 |
| Vipingo | 1/1 (100%) | 0/1 (0%) | 1 |
| **Total** | **74/299 (24.8%)** | **17/299 (5.7%)** | **299** |

<sup>1</sup> Village where one participant was found with circulating *Plasmodium ovale* parasites. Shown in parenthesis are percent positive per village.

**Figure 2.**
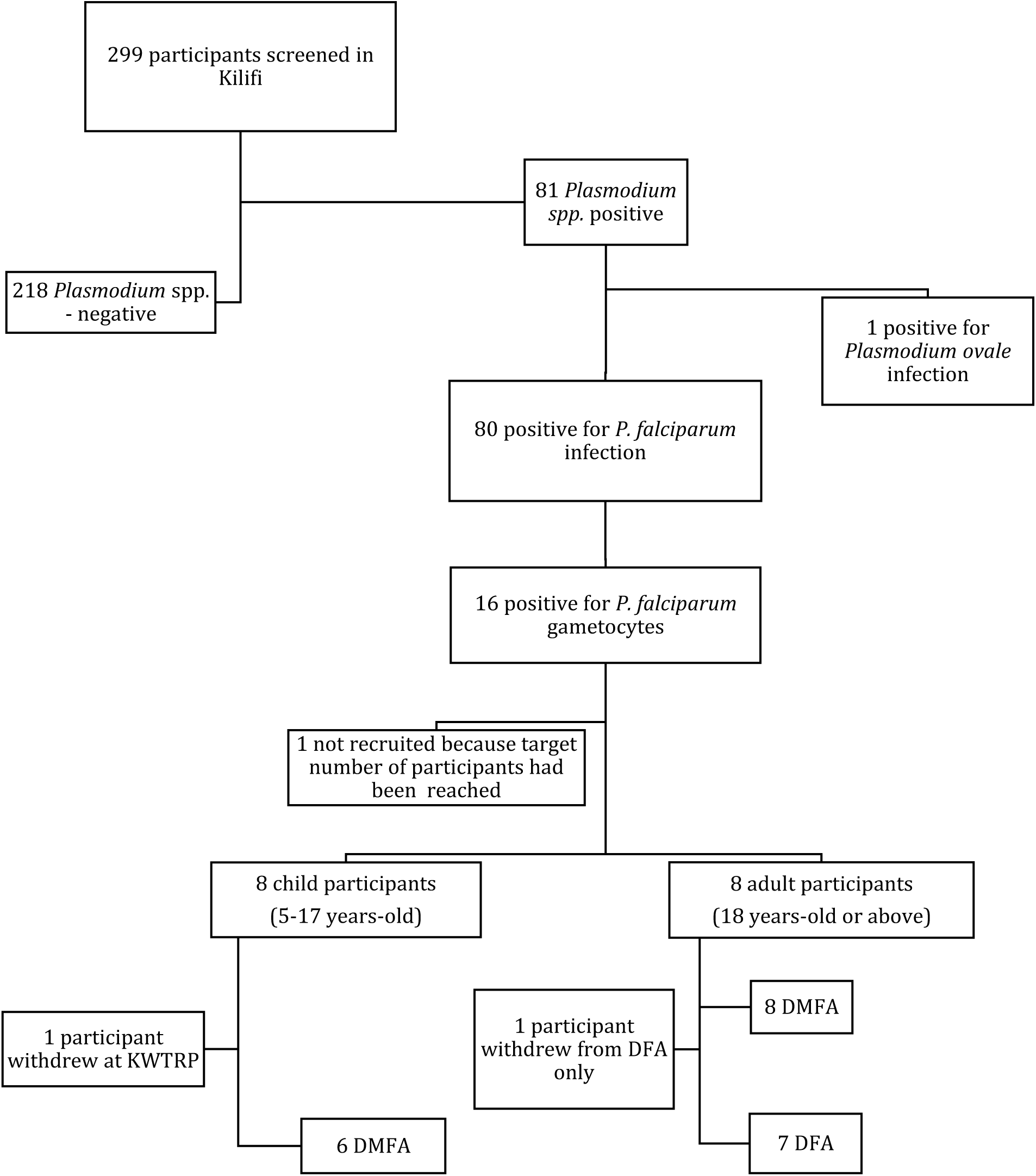
Flow chart showing number of adult and child participants that were screened, and recruited for direct membrane feeding assay (DMFA) and direct feeding assay (DFA).

Parasite prevalence decreased with age (Figure 3) (Table 2). Older adults (40-49-year-old) were the least likely to be found with circulating parasites, only 5% (2/44) had asexual stage parasites and 2% (1/44) sexual stage parasite in circulation. The highest parasite prevalence was found in children 5-9 years old; 36% (16/45) of the children screened were positive for asexual stage parasites, and 11% (5/45) were positive for gametocytes. Older children and adolescents (10-19 years old) had only slightly lower parasite prevalence than the younger age group, 32% (28/87) of the children were found with asexual parasites and 6% (5/87) with circulating gametocytes. Young (20-29 years old) and middle-aged adults (30-39 years old) had comparable parasite prevalence with respectively 24% (16/66) and 21% (12/57) found with asexual parasite stages, and 6% (4/66) and 4% (2/57) found with sexual parasite stages.

**Table 2.** Number of participants by age group screened for asexual (A) and sexual stage (B) *Plasmodium spp*. parasites. Median parasitemia per mcl and interquartile range (based on number of parasites counted per 200 WBC assuming 8000 WBC/mcl), minimum and maximum parasites per mcl and number of participants recruited for direct membrane feeding assay (DMFA) and direct feeding assay (DFA).

| **A – Asexual parasites** |  |  |  |  |  |  |  |
| --- | --- | --- | --- | --- | --- | --- | --- |
| **Age** | No. participants<sup>1</sup> | Asexual parasite prevalence | Median parasites / mcl (IQR) | Min - Max / mcl |  |  |  |
| **5-9** | 16/45 | 0.36 | 600 (160 – 1'640) | 80 – 31'040 |  |  |  |
| **10-19** | 28/87 | 0.32 | 280 (120 - 940) | 80 – 26'560 |  |  |  |
| **20-29** | 16/66 | 0.24 | 280 (140 – 1'140) | 40 – 9'160 |  |  |  |
| **30-39** | 12/57 | 0.21 | 220 (140 – 700) | 80 – 5'760 |  |  |  |
| **40-49** | 2/44 | 0.05 | 200 (120 – 280) | 120 - 280 |  |  |  |
| **B – Gametocytes** |  |  |  |  |  |  |  |
| **Age** | No. participants | Gametocyte prevalence | Median parasites / mcl (IQR) | Min - Max / mcl | No. participants recruited | DMFA | DFA |
| **5-9** | 5/45 | 0.11 | 280 (160 – 280) | 40 – 1'240 | 2<sup>2,3,4</sup> | 2 | 0 |
| **10-19** | 5/87 | 0.06 | 80 (80 – 80) | 40 - 80 | 4 | 4 | 1<sup>5</sup> |
| **20-29** | 4/66 | 0.06 | 300 (240 – 360) | 200 - 400 | 4 | 4 | 4 |
| **30-39** | 2/57 | 0.04 | 520 (80 – 960) | 80 - 960 | 2 | 2 | 1<sup>6</sup> |
| **40-49** | 1/44 | 0.02 | 160 (160 – 160) | 160 - 160 | 1 | 1 | 1 |

<sup>1</sup> Number of participants found with circulating parasites/total participants screened
<sup>2</sup> One individual withdrew participation out of fear of needles.
<sup>3</sup> One gametocyte-positive individual had a *Plasmodium. ovale* infection and so did not meet the study's inclusion criteria.
<sup>4</sup> One individual was randomly excluded because the target number of 14 participants had been reached.
<sup>5</sup> DFA was performed on a 19-year old volunteer.
<sup>6</sup> One of the adult participants consented to DMFA but not to DFA for fear of witchcraft.

**Figure 3.**
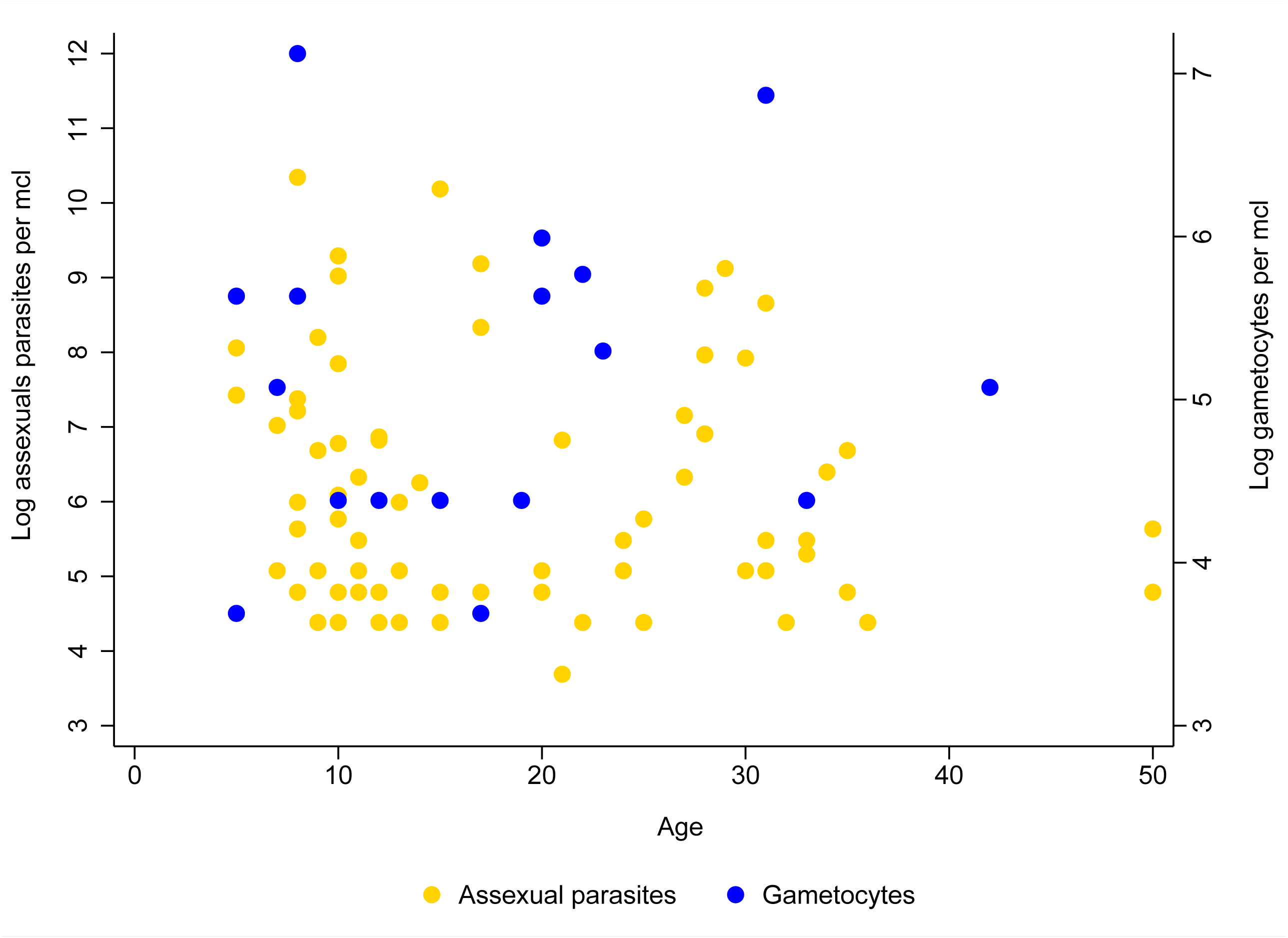
Log transformed asexual (left y axis) and sexual parasites (right y axis) per mcl by age group (x axis) from the parasitological survey done in Kilifi South, Kenya between 28th May and 18^th^ September 2018.

Asexual parasite densities per mcl was highest in the children 5-9 years old [Median=600; IQR= (160-1640], with highest parasite densities recorded in an 8-year-old child (31’040 parasites/mcl). Older children and young adults had lower median parasite densities compared to the younger children, whilst older adults (40-49 years old) had lowest parasite densities (Median=200; IQR= (120-280)].

Fifteen gametocyte carriers, eight adults and seven children, were recruited and invited to participate in mosquito feeding assays. One child withdrew before sample could be collected due to fear of needles, and one adult participant consented to the blood draw for DMFA but withdrew from participating in the direct feeding assay because of beliefs in witchcraft. DMFA was performed with blood of all 14 participants, and DFA was performed on 7 of the adult participants (Table 2).

### Infectivity to mosquitoes

Of the fourteen asymptomatic gametocyte carriers that were recruited for mosquito feeding assays, 9 resulted in successful sporozoite infections confirmed by CSP ELISA (Table 3). High gametocyte densities did not necessarily result in high sporozoite positivity rate (Figure 4). For instance, the highest mosquito infection rate (47.0% infection by DFA and 17.6% by DMFA) resulted from a participant with low gametocyte densities (80 gametocytes/mcl). Concurrent DMFA and DFA were performed on 7 adult participants, of which 4 participants were able to infect mosquitoes. Unfortunately, mosquitoes fed by DFA from one participant escaped and had to be killed before reaching 14-days post feeding to maintain biosecurity. Overall mosquito infection rate by DMFA was 4.9%. Concurrent DMFA and DFA, performed on the same participant, resulted in 11.5% and 23.7% infection rates, respectively.

**Table 3.** Infectivity of individual participants based on feeding assay and positivity by CSP-ELISA

| Participant | Age | Gametocytes per mcl | DMFA<sup>1</sup><br>% (n/N) | DFA<sup>1</sup><br>% (n/N) | Total<sup>1</sup><br>% (n/N) |
| --- | --- | --- | --- | --- | --- |
| Adult 1 | 42 | 160 | 0.0% (0/80) | 12.5% (3/24) | 2.9% (3/104) |
| Adult 2 | 23 | 200 | 1.4% (1/74) | 7.1% (2/28) | 2.9% (3/102) |
| Adult 3 | 19 | 80 | 17.6% (15/85) | 47.0% (8/17) | 22.6% (23/102) |
| Adult 4<sup>2</sup> | 20 | 400 | 26.9% (66/245) | 28.0% (7/25) | 27% (73/270) |
| Adult 5<sup>3</sup> | 31 | 960 | 17.0% (26/153) | - | 17.0% (26/153) |
| Adult 6 | 33 | 80 | 0.7% (1/141) | - | 0.7% (1/141) |
| Adult 7 | 22 | 320 | 0% (0/61) | 0% (0/15) | 0% (0/76) |
| Adult 8 | 20 | 280 | 0% (0/137) | 0% (0/33) | 0% (0/170) |
| Child 1 | 10 | 80 | 2.1% (2/94) | - | 2.1% (2/94) |
| Child 2 | 5 | 40 | 0.5% (1/199) | - | 0.5% (1/199) |
| Child 3 | 17 | 40 | 2.6% (2/77) | - | 2.6% (2/77) |
| Child 4 | 7 | 160 | 0% (0/240) | - | 0% (0/240) |
| Child 5 | 15 | 80 | 0% (0/66) | - | 0% (0/66) |
| Child 6 | 12 | 80 | 0% (0/87) | - | 0% (0/87) |
| **Total** |  |  | **4.9% (114/1739)** | **15.8% (20/142)** | **7.1% (134/1881)** |

<sup>1</sup> Percent positive of mosquitoes by CSP-ELISA based on mosquito feeding assay
<sup>2</sup> All mosquitoes fed by DFA on this participant were killed before reaching 14 days-old to maintain biosecurity.
<sup>3</sup> Withdrew participation from DFA over fear of witchcraft. *Leave-one-out cross validations (LOOCV)*

**Figure 4.**
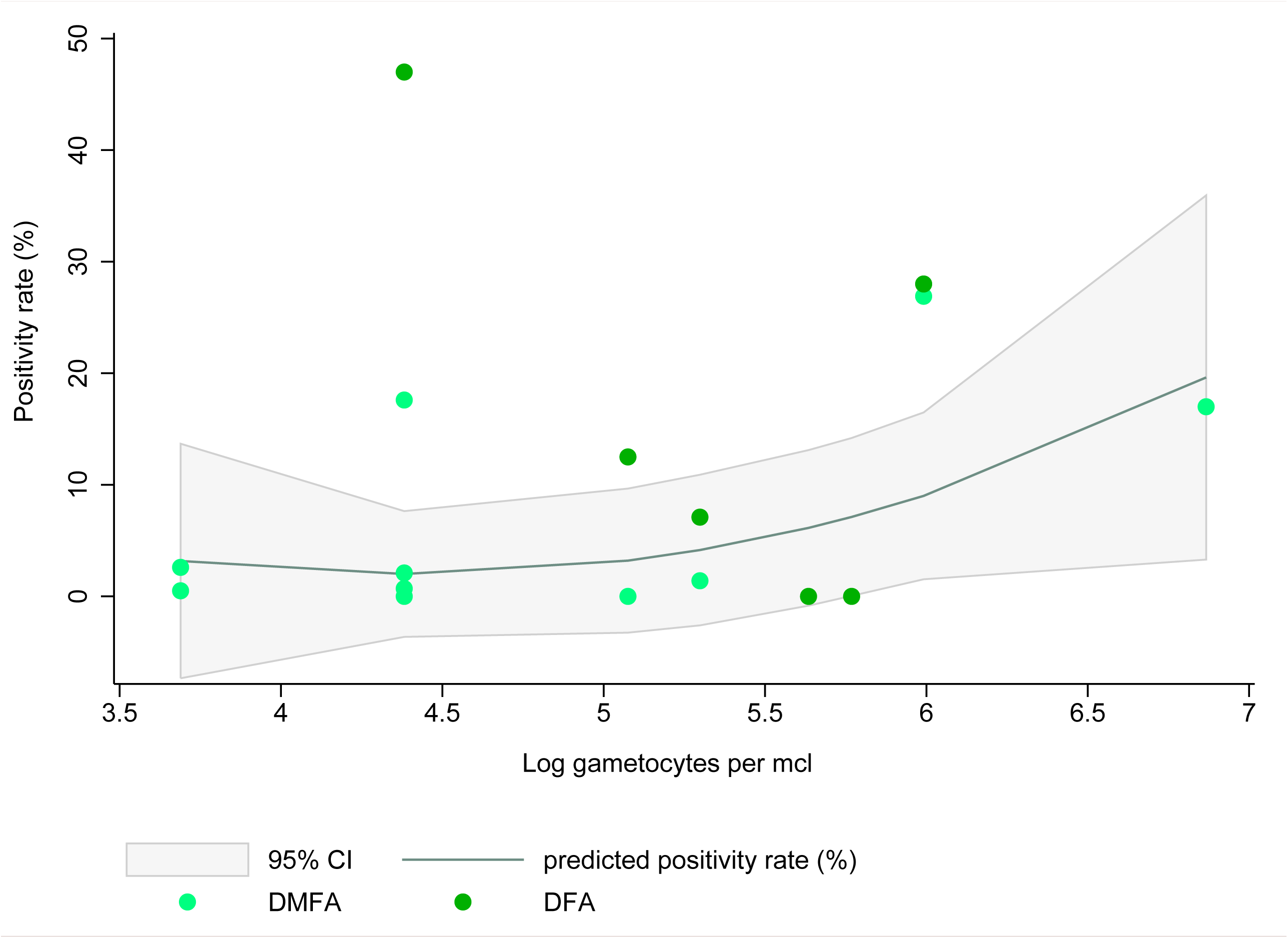
Log transformed gametocytes per mcl (x axis) and percentage of mosquitoes that became sporozoite positive (positivity rate) (y axis) after DMFA and DFA. Fitted predicted positivity rate and 95% confidence interval (CI).

Overall, 1,881 individual NIR absorbance spectra (500-2400 nm) from potential sporozoite-infected mosquitoes were taken, of which 1739 mosquitoes had been fed via DMFA and 142 via DFA. A total of 134 NIRS absorbance spectra corresponding to sporozoite-infected mosquitoes were obtained. Three training datasets were analyzed using PLS through LOOCV on Grams Plus/ IQ software for the development of a predictive model to distinguish between sporozoite-infected and uninfected mosquitoes.

In the first analysis, a pool of 134 spectra from sporozoite-infected mosquitoes were randomly divided in two, half of the spectra were assigned to a training dataset (67) and the other half to a test dataset (67) to be used for validating the model if it showed good self-predictive ability. Each spectrum from a sporozoite-infected mosquito was matched with a spectrum corresponding to an uninfected mosquito from the same experimental feed. Matching was done to reduce confounding effects such as parasite load and participant immunity as the assays were done with whole blood. Thus, a total of 67 spectra corresponding to sporozoite-infected mosquitoes were matched with 67 spectra from uninfected mosquitoes (134 spectra: 67 uninfected/67 sporozoite-infected). The LOOCV showed poor model predictive ability, (R2= 0.029). the variation between sporozoite-infected spectra and uninfected spectra could not be explained by the mosquito’s infectious status (see Analysis 1 in supplementary file).

A second analysis was performed to ascertain that the two NIR spectra categories were not distinct by increasing the number of spectra in the training dataset. A second training dataset included all the spectra from sporozoite-infected mosquitoes (134) and was matched with respective uninfected mosquitoes (134 sporozoite-infected/134 uninfected). A second LOOCV was performed using the larger training dataset (268 spectra: 134 uninfected/134 sporozoite-infected). The model was not improved (R2 = 0.00000009) and remained unable to differentiate the spectra from the two mosquito groups (see analysis 2 in supplementary file).

Uninfected mosquitoes in the first and second analysis had been matched to infected ones from the same experimental feed, consequently it was possible that the uninfected mosquitoes had been exposed to infectious gametocytes but had not developed sporozoite infection. Hence, a third analysis was performed to rule out the possibility that an arthropod immune response had affected the NIR spectra profiles of the uninfected mosquitoes. A third training dataset was composed of 99 sporozoite-infected mosquitoes from the three participants with highest experimental infection rates (Adult 1, 3, 4), and 99 uninfected mosquitoes from three unsuccessful experiments (i.e. mosquito feeding experiments that did not lead to at least one infected mosquito). An LOOCV was performed to give information on the prediction ability of the model using a training dataset composed of 198 spectra (99 sporozoite-infected/99 uninfected from unsuccessful experiments). Yet again, the PLS model was unable to distinguish between the two spectra groups (R2= 0.049). (see Analysis 3 in supplementary file).

A calibration could not be generated based on any of the training datasets as the variation between the two groups of spectra (sporozoite-infected mosquitoes Vs uninfected mosquitoes) could not be explained by any of the models using PLS.

## DISCUSSION

The present study shows that mosquito transmission studies using DMFA and DFA can be readily conducted in Kilifi by targeting an area known as a malaria hotspot. Screening was targeted to areas where malaria cases had recently been reported at the health care facility which may have overestimated the prevalence of malaria in the area compared to other published studies from the same area [13]. The targeted screening approach proved to be a successful strategy for identifying gametocyte carriers in the community.

The study achieved 9 out of 14 successful experiments, defined as at least one mosquito becoming infected after blood feeding based on the detection of sporozoites in each individual mosquito by ELISA. The highest infection rate resulted from a participant who had relatively low gametocytemia (Table 3). It is known that mosquito infectivity to gametocytes is dependent of multiple factors asides from gametocyte density, including the adequate ratio of male to female parasites, their level of maturity as well as mosquito microbiome and immunity [14]. In the present experiment, mosquitoes were all from the same strain and reared in the same insectary conditions therefore had plausible similar microbiome and immunity.

DFA consistently produced higher infection rates. This is in line with other studies which have compared the two methods [9, 10]. Gametocytes are known to sequester in the microvasculature, thus venous blood used for DMFA may have contained fewer parasites compared to capillary blood. It is also possible that DFA was more successful because unlike DMFA the method does not require blood handling during venipuncture which may lead to exflagellation of the male gametocytes and loss of infectiousness.

The three training datasets used to investigate if a PLS predictive model could be generated to predict sporozoite infections status in mosquitoes demonstrated poor out-of-sample predictive accuracy. The models were unable to satisfactorily predict the left-out spectra *(k-1)*. The PRESS showed no latent factor could be identified to explain the variation between spectra groups (sporozoite-infected mosquitoes Vs uninfected mosquitoes) (see supplementary file). R^2^ and predicted R^2^ values for three models evaluated were very low indicating its inadequacy to explain the variation between the constituents. Given the poor self-explanatory ability of the three PLS models, the present study was unable to generate a calibration for distinguishing sporozoite infection status of *A. gambiae* s.s. mosquitoes.

The present findings are at variance with a previous study which showed that PLS models based on the NIR absorbance spectra profile, could accurately predict both oocyst and sporozoite infection status of *A. gambiae* s.s. mosquitoes infected with laboratory-cultured *P. falciparum* (PfN54) parasites [4]. Differences between the two studies may explain the conflicting results.

In the previous study, mosquitoes were infected with lab-reared gametocytes through standard membrane feeding assays (SMFA) using higher parasite concentrations than what is usually found in a human gametocyte carrier. It is likely that mosquitoes in the present study were, in comparison, exposed to a much smaller number of parasites which more closely represents natural infections in nature Findings from the previous study also concluded that the prediction accuracy of the model dropped when mosquitoes had reduced infection loads (measured as number of parasite genomes). A limitation of the current study is that parasite quantification in the mosquito was not performed through qPCR (quantitative polymerase chain reaction). ELISA was chosen as reference test as it is the assay of choice for field entomologists to identify sporozoite-infected mosquitoes and measure sporozoite rates.

The training datasets of Analysis 1 and 2 included spectra of uninfected mosquitoes that had been matched with spectra of sporozoite-infected mosquitoes fed on the same infectious blood meal. These uninfected mosquitoes were exposed to an infectious blood meal but had not developed sporozoites. It is possible that during feeding these mosquitoes did not pick-up a male and female gametocyte for subsequent differentiation into gametes and mating inside the mosquito gut, or alternatively they may have blocked infection through an immune response. In comparison, the predictive model of the previous study [4] was characterized by uninfected mosquitoes that had been fed neutralized gametocytes, which are unlikely to trigger an immune response in the arthropod vector. To evaluate if the mosquito’s immune response had affected the NIR profile of the uninfected mosquitoes Analysis 3 was done whereby spectra were no longer matched by experimental feed. In Analysis 3, spectra from sporozoite-infected mosquitoes were compared to spectra from uninfected mosquitoes fed during unsuccessful experiments (i.e. an experimental feed where none of the fed mosquitoes developed sporozoite infection). Yet again, the predictive ability of the model remained poor and it is unlikely that arthropod immune response significantly affected the NIR spectra profile of the mosquitoes.

PLS regression is the most widely used approach for developing predictive models to distinguish between entomological parameters such as mosquito age, species as well as *Wolbachia* infection [6, 7, 15]. Other machine learning methods, such as artificial neural networks have been recently suggested as alternative analytical approaches [16, 17] and it is possible that these may improve the prediction ability of models to distinguish between NIR spectra of infected and uninfected mosquitoes.

## CONCLUSIONS

While the findings presented here were unable to reproduce and replicate previous finding [4],thes results are important as the will inform future studies seeking novel high-throughput methods for parasite detection in mosquitoes. The results of this study were not consistent with previous published research on NIRS for detection of sporozoite infection in the same mosquito species thought different strains, and this may be explained by the context of the experiments. Further research may clarify the discrepancy in the findings. It is essential that new technologies for identifying infections in mosquitoes are assessed and developed in the field setting at an early stage as this is their primary application.

## LIST OF ABBREVIATIONS

CSP ELISA: Circumsporozoite Enzyme Linked Immunosorbent Assay
DMFA: Direct Membrane Feeding Assay
DFA: Direct Feeding Assay
KWTRP: Kenya Medical Research Institute, Wellcome Trust Research Programme
KOPD: Kilifi Out-patient Department
NIRS: Near infrared spectroscopy
LOOCV: Leave-one-out Cross Validations
PLS: Partial Least Squares
WBC: White Blood Cells

## ETHICS APPROVAL

Individual consent was obtained before each screening, parental consent was obtained for minors (5-17 years old) and assent was obtained for children between 12 and 17 years old. Ethical approval (KEMRI/SERU/CGMR-C/082/3523) was obtained from SERU-KEMRI (Scientific and Ethics Review Unit of the Kenya Medical Research Institute).

## CONSENT FOR PUBLICATION

Not applicable.

## AVAILABILITY OF DATA AND MATERIAL

All the data necessary to interpret and replicate the finding on this paper have been made publicly available on the data repository Harvard dataverse [18]. This includes anonymized screening results, as well as DMFA and DFA assay results; NIR spectra of all the specimens (spc files) with specification to whether they correspond to sporozoite-infected or uninfected mosquitoes.

## AUTHORS CONTRIBUTIONS

MFM drafted the manuscript and designed the experiment with input from PB, FD and MCK. MFM led the study, including obtaining study ethics approvals, participant screening and recruitment, mosquito feeding assays, data collection and analysis. FM and JK performed the CSP ELISA assays. MFM and MW scanned the mosquitoes and collected the NIR data. JW managed the team of field workers who collected the blood smears for screening. RM read all the study blood smears. MM and FM maintained the insectary and prepared mosquitoes for the experimental assays. MH advised on all clinical aspects of the study and ensured clinical care was provided. All authors read and commented on drafts of the manuscript and approved the final version for publication.

## ACKNOWLEDGEMENTS

We would like to thank the medical officers Joseph Weya and Kenneth Kiogora for attending to the participants at the KOPD; the KWTRP field worker team; Christopher Nyundo for assisting with a map of the study site; the KEMRI community representatives; and the KEMRI community liaison group (CLG). Mostly, the authors thank the communities of Junju and Pingilikani for their participation. In addition, we acknowledge, the following reagent was obtained through BEI Resources, NIAID, NIH: *Plasmodium falciparum* Sporozoite ELISA Reagent Kit, MRA-890, contributed by Robert A. Wirtz

## ADDITIONAL FILES

Additional file 1.docx: GRAMS output figures and table describing the mosquitoes used in the three training datasets by participant and infection status (determined by Csp-ELISA).

## REFERENCES

1. WHO: World Malaria Report. World Health Organization: Geneva, Switzerland; 2019.

2. malERA: An updated research agenda for insecticide and drug resistance in malaria elimination and eradication. PLoS Med 2017, 14:e1002450.

3. Burkot TR, Zavala F, Gwadz RW, Collins FH, Nussenzweig RS, Roberts DR: Identification of malaria-infected mosquitoes by a two-site enzyme-linked immunosorbent assay. Am J Trop Med Hyg 1984, 33:227–231.

4. Maia MF, Kapulu M, Muthui M, Wagah MG, Ferguson HM, Dowell FE, Baldini F, Ranford-Cartwright L: Detection of Plasmodium falciparum infected Anopheles gambiae using near-infrared spectroscopy. Malar Jl 2019, 18:85.

5. Tusting LS, Bousema T, Smith DL, Drakeley C: Measuring changes in Plasmodium falciparum transmission: precision, accuracy and costs of metrics. Adv Parasitol 2014, 84:151–208.

6. Mayagaya VS, Michel K, Benedict MQ, Killeen GF, Wirtz RA, Ferguson HM, Dowell FE: Non-destructive determination of age and species of Anopheles gambiae s.l. using near-infrared spectroscopy. Am J Trop Med Hyg 2009, 81:622–630.

7. Sikulu M, Killeen GF, Hugo LE, Ryan PA, Dowell KM, Wirtz RA, Moore SJ, Dowell FE: Near-infrared spectroscopy as a complementary age grading and species identification tool for African malaria vectors. Parasit Vectors 2010, 3:49.

8. Esperança PM, Blagborough AM, Da DF, Dowell FE, Churcher TS: Detection of Plasmodium berghei infected Anopheles stephensi using near-infrared spectroscopy. Parasit Vectors 2018, 11:377.

9. Bousema T, Dinglasan RR, Morlais I, Gouagna LC, van Warmerdam T, Awono-Ambene PH, Bonnet S, Diallo M, Coulibaly M, Tchuinkam T, et al: Mosquito feeding assays to determine the infectiousness of naturally infected Plasmodium falciparum gametocyte carriers. PLoS ONE 2012, 7:e42821.

10. Diallo M, Toure AM, Traore SF, Niare O, Kassambara L, Konare A, Coulibaly M, Bagayogo M, Beier JC, Sakai RK, et al: Evaluation and optimization of membrane feeding compared to direct feeding as an assay for infectivity. Malar J 2008, 7:248.

11. Kenya MoH: National Guidelines for the diagnosis, treatment and prevention of malaria in Kenya. 5th Edition edition. Nairobi, Kenya: National Malaria Control Program; 2016.

12. Beier MS, Schwartz IK, Beier JC, Perkins PV, Onyango F, Koros JK, Campbell GH, Andrysiak PM, Brandling-Bennett AD: Identification of malaria species by ELISA in sporozoite and oocyst infected Anopheles from western Kenya. Am J Trop Med Hyg 1988, 39:323–327.

13. Muthui MK, Mogeni P, Mwai K, Nyundo C, Macharia A, Williams TN, Nyangweso G, Wambua J, Mwanga D, Marsh K, et al: Gametocyte carriage in an era of changing malaria epidemiology: A 19-year analysis of a malaria longitudinal cohort. Wellcome Open Res 2019, 4.

14. Sinden RE, Blagborough AM, Churcher T, Ramakrishnan C, Biswas S, Delves MJ: The design and interpretation of laboratory assays measuring mosquito transmission of Plasmodium. Trends Parasitol 2012, 28:457–465.

15. Sikulu-Lord MT, Maia MF, Milali MP, Henry M, Mkandawile G, Kho EA, Wirtz RA, Hugo LE, Dowell FE, Devine GJ: Rapid and Non-destructive Detection and Identification of Two Strains of Wolbachia in Aedes aegypti by Near-Infrared Spectroscopy. PLoS Negl Trop Dis 2016, 10:e0004759.

16. Milali MP, Sikulu-Lord MT, Kiware SS, Dowell FE, Corliss GF, Povinelli RJ: Age grading An. gambiae and An. arabiensis using near infrared spectra and artificial neural networks. PLoS One 2019, 14:e0209451.

17. Milali MP, Kiware SS, Govella NJ, Okumu F, Bansal N, Bozdag S, Charlwood JD, Maia MF, Ogoma SB, Dowell FE, et al: An autoencoder and artificial neural network-based method to estimate parity status of wild mosquitoes from near-infrared spectra. PLoS One 2020, 15.

18. Maia MF, Wagah MG, Karisa J, Mure F, Mwakesi R, Muturi M, Wambua J, Hamaluba M, Dowell F, Bejon P, Kapulu MC: Data for: Evaluation of near infrared spectroscopy for sporozoite detection in mosquitoes infected with wild-strain parasites from asymptomatic gametocyte carriers in Kilifi Kenya. https://doi.org/10.7910/DVN/IVFTLB, Harvard Dataverse, V1. 2020.

