## Supplementary information for "Evaluation of near infrared spectroscopy for sporozoite detection in mosquitoes infected with wild-strain parasites from asymptomatic gametocyte carriers in Kilifi Kenya"

---

**Maia et al. 2020**

---

*Introduction to supplementary information*

A predictive model for distinguishing between NIR absorbance spectra of sporozoite-infected and uninfected *Anopheles gambiae* s.s was explored using partial least square (PLS) regression. Spectra were added to Grams Plus/IQ (Thermo Galactic, Salem, NH, USA). Mosquitoes of same age were infected with wild *Plasmodium falciparum* by direct membrane feeding assay (DMFA) and direct feeding assay (DFA) as explained in the main manuscript.

**Analysis 1**

The pool of 134 spectra of sporozoite-infected mosquitoes was first randomly divided in two and assigned to either a training or test dataset (Table 1). All the spectra from infected mosquitoes were matched with spectra of uninfected mosquitoes from the same feeding experiment and assay. Thus each dataset (training and test) was composed of 67 spectra from sporozoite-infected and 67 spectra from uninfected counterparts. A leave-one-out cross validation using PLS regression was done to evaluate NIR spectral differences between sporozoite-infected and uninfected *Anopheles gambiae* s.s.

**Supplementary Table 1** – Number of spectra from uninfected and sporozoite-infected mosquitoes, determined by CSP ELISA selected for inclusion in the training and test datasets of Analysis 1.

| | **Training dataset** | | **Test dataset** | |
|---|---|---|---|---|
| | **CSP-Positive** | **Negative** | **CSP-Positive** | **Negative** |
| Adult 1 | 2 | 2 | 1 | 1 |
| Adult 2 | 1 | 1 | 2 | 2 |
| Adult 3 | 12 | 12 | 11 | 11 |
| Adult 4 | 36 | 36 | 37 | 37 |
| Adult 5 | 13 | 13 | 13 | 13 |
| Adult 6 | 1 | 1 | 0 | 0 |
| Child 1 | 1 | 1 | 1 | 1 |
| Child 2 | 0 | 0 | 1 | 1 |
| Child 3 | 1 | 1 | 1 | 1 |
| **TOTAL** | **67** | **67** | **67** | **67** |

All .asd files were converted to .spc uploaded onto Grams Plus/IQ and given either a constituent of 1 (spectra from uninfected mosquitoes) or a constituent of 2 (spectra of sporozoite-infected mosquitoes). The spectra were observed and any obvious outliers were removed from the analysis. Spectra ID SJA019201200 (corresponding to sporozoite-infected) was excluded because it was considered too flat.

*Results*

The model's predictive ability was poor (**Supplementary** Figures 1 and 2). The variation between the spectra could not be explained by their infection status ($R^2$= 0.029). The test dataset was not queried given the poor prediction ability of the training dataset.

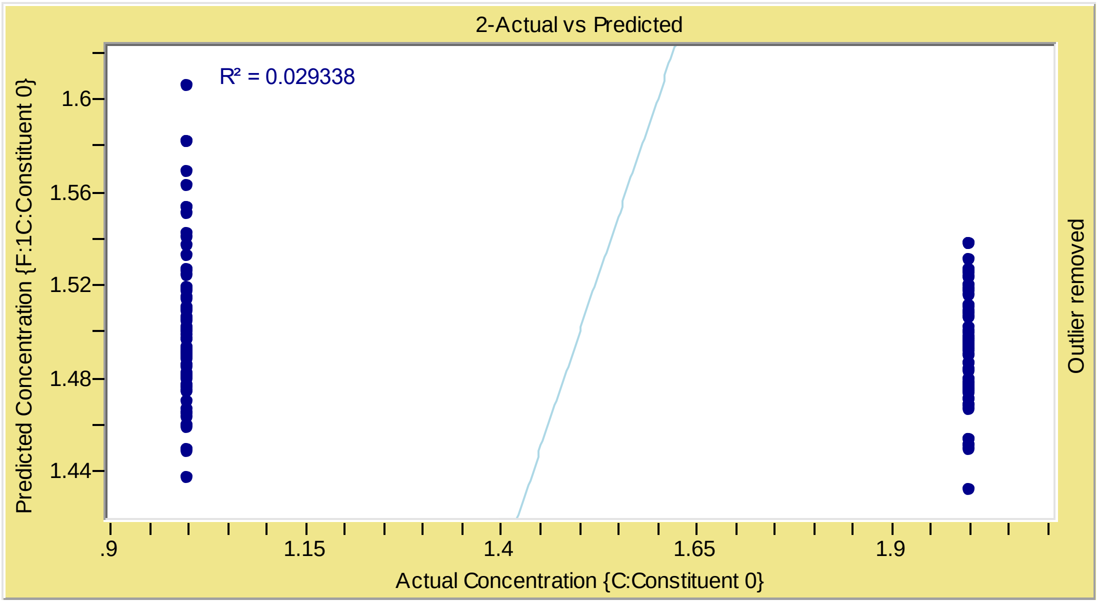


**Supplementary Figure 1** – Actual Vs Predicted plot. $R^2 = 0.029$. Constituent 1: uninfected; constituent 2: Sporozoite-infected (Analysis 1).

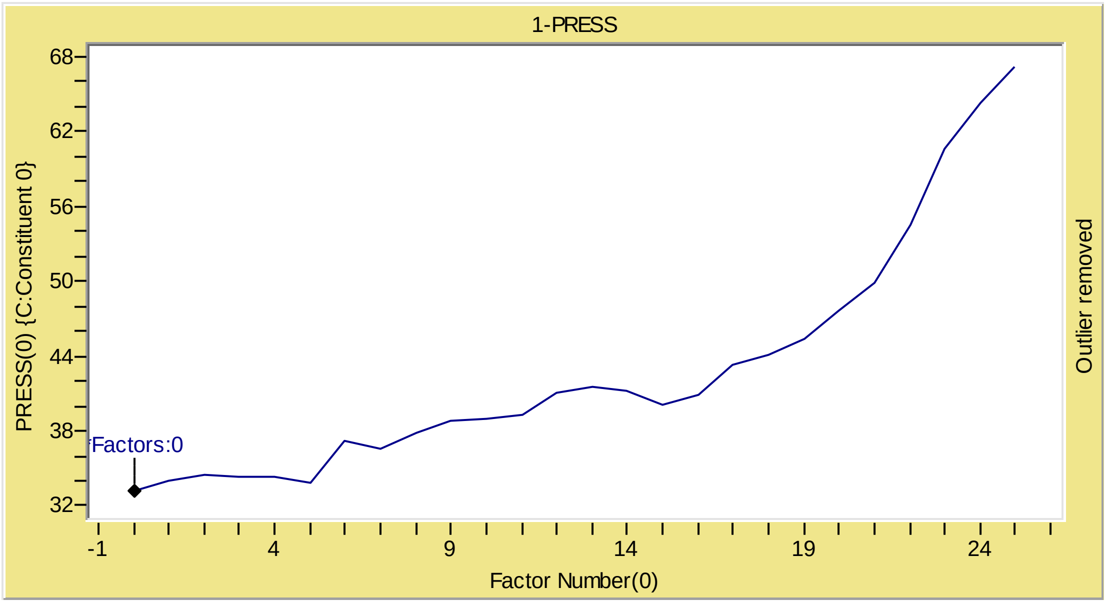


**Supplementary Figure 2 -** Prediction residual sum of squares (PRESS) curve (Analysis 1). No latent factor could be extracted to form a predictive model.

**Analysis 2**

Given the poor prediction ability of the model presented in Analysis 1 the authors attempted to improve the model by increasing the number of samples in the training dataset. A training dataset composed of 266 spectra were added to Grams IQ for evaluating differences in NIR absorbance spectra between infected and uninfected *Anopheles gambiae* s.s. mosquitoes (Table 2). Mosquitoes were infected with wild *Plasmodium falciparum* by DMFA and DFA as explained in the main manuscript. Special care was taken to ensure all infected mosquitoes were matched in the database with uninfected mosquitoes from the same feeding assay.

**Supplementary Table 2** – Number of spectra from uninfected and sporozoite-infected mosquitoes, determined by CSP-ELISA selected for inclusion in the training dataset of Analysis 2.

| | **Training dataset** | |
|---|---|---|
| | **CSP-Positive** | **Negative** |
| Adult 1 | 3 | 0 |
| Adult 2 | 3 | 3 |
| Adult 3 | 23 | 23 |
| Adult 4 | 73 | 73 |
| Adult 5 | 26 | 26 |
| Adult 6 | 1 | 1 |
| Child 1 | 2 | 2 |
| Child 2 | 1 | 1 |
| Child 3 | 2 | 2 |
| **TOTAL** | **134** | **134** |

Similar to previous analysis, spectra were assigned either a constituent of 1 (uninfected mosquitoes) or a constituent of 2 (sporozoite-infected mosquitoes). The spectra were observed and any obvious outliers were removed from the analysis. Spectra ID SJA019201200 (corresponding to sporozoite-infected) was again excluded because it was considered too flat.

*Results*

The models predictive ability was not improved (Figure 3 and 4). The test dataset was not queried given the poor prediction ability of the training dataset.

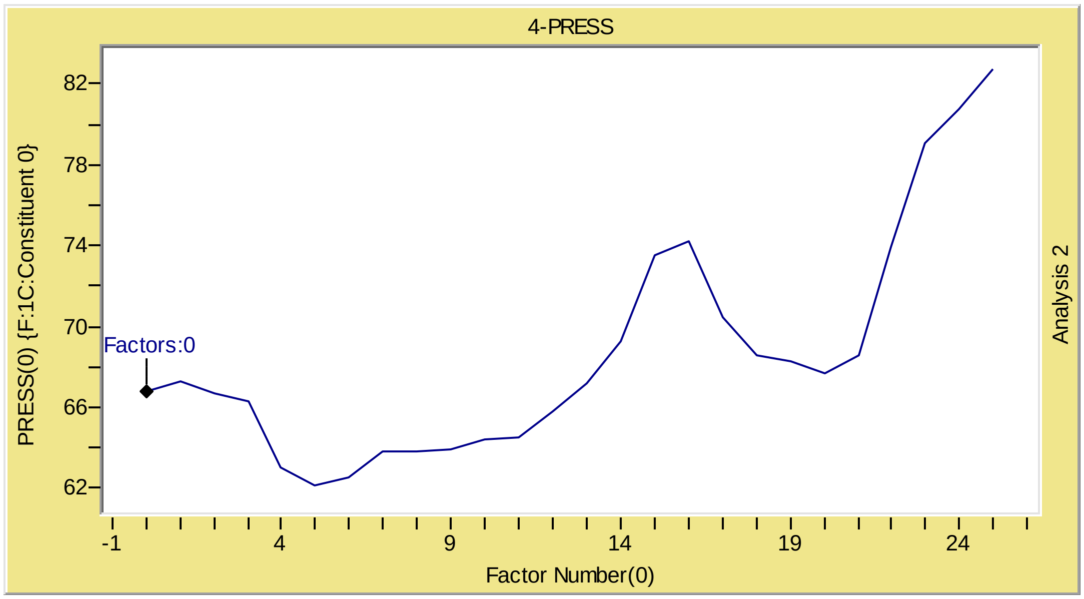


**Supplementary Figure 3 -** Prediction residual sum of squares (PRESS) curve (Analysis 2). No latent factor could be extracted to form a predictive model.

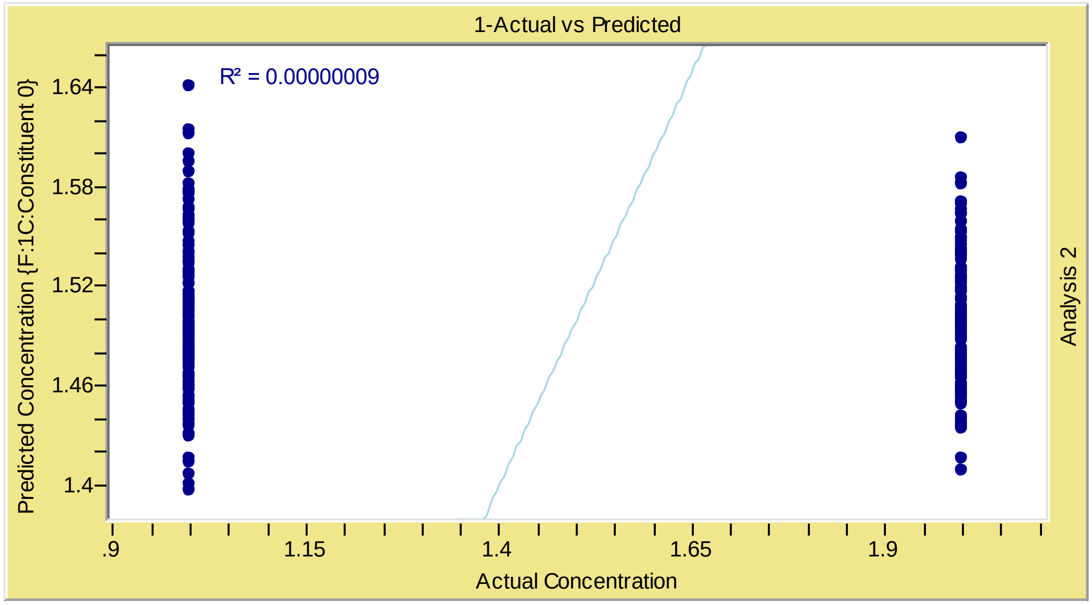


**Supplementary Figure 4** – Actual Vs Predicted plot. $R^2 = 0.00000009$. Constituent 1: uninfected; constituent 2: Sporozoite-infected (Analysis 2).

**Analysis 3**

The predictive models developed in analysis 1 and 2 were both unable to distinguish between the NIR absorbance spectra of the sporozoite-infected and uninfected mosquitoes included in the respective training datasets. The training datasets of Analysis 1 and 2 were composed of infected and matched uninfected spectra from the same experiment and assay. A third analysis was done to rule out if the arthropod's immune response to the parasites in the infectious blood had affected its NIR spectra profile. This was done by including spectra from an unsuccessful experiment i.e. an experiment whereas neither DMFA nor DFA resulted in at least one infected mosquito. This was done under the assumption that these mosquitoes would have not become infected because they did not ingest infectious gametocytes rather than due to their immune response. A training dataset was randomly selected to test this hypothesis composed of 99 sporozoite-infected mosquitoes and 99 uninfected mosquitoes from unsuccessful experiments. Consequently, in this model matching of spectra by experimental assay was not done.

**Supplementary Table 3** – Number of spectra from uninfected and sporozoite-infected mosquitoes, determined by CSP ELISA selected for inclusion in the training dataset of Analysis 3.

| | **Training dataset** | |
|---|---|---|
| | **Csp-Positive** | **Negative** |
| Adult 1 | 3 | 0 |
| Adult 3 | 23 | 0 |
| Adult 4 | 73 | 0 |
| Child 4[1] | 0 | 33 |
| Child 5[1] | 0 | 33 |
| Child 6[1] | 0 | 33 |
| **TOTAL** | **99** | **99** |

[1]Spectra corresponding to mosquitoes from unsuccessful mosquito feeding assays.

Constituents remained the same (1: uninfected; 2: infected).

*Results*

The models predictive ability was not improved after limiting the uninfected spectra of the training dataset to spectra from unsuccessful transmission assays (Figure 5 and 6). Assuming that mosquitoes from unsuccessful transmission assays were not exposed to infectious parasites, it is unlikely that the mosquito's immune response was affecting the NIR profiles. The current model could not explain the variance between spectra infected and uninfected (likely unexposed) mosquitoes ($R^2$= 0.049). The test dataset was not queried given the poor prediction ability of the training dataset.

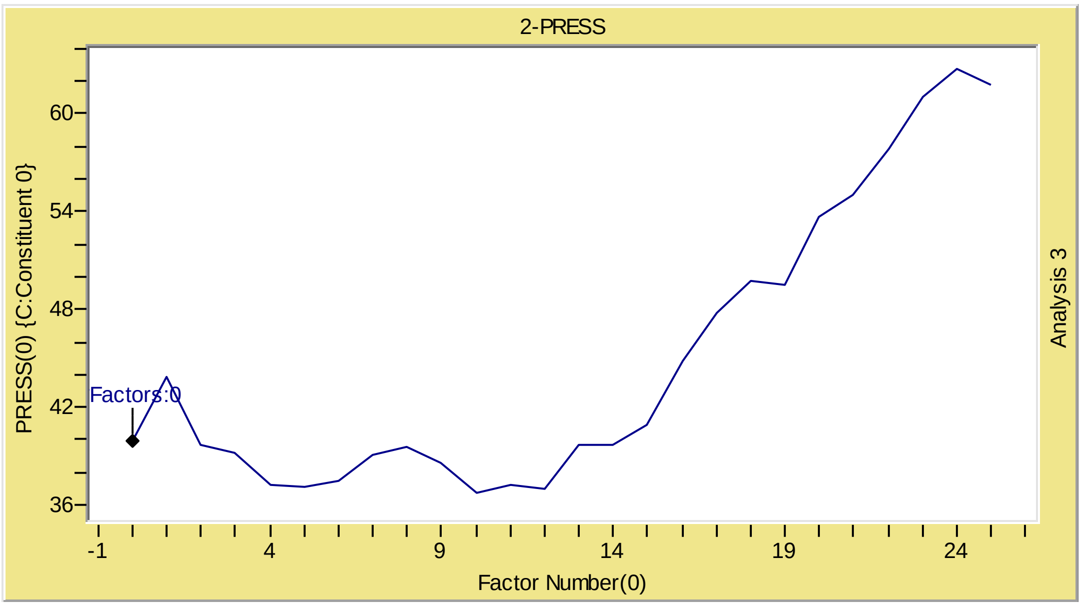


**Supplementary Figure 5 -** Prediction residual sum of squares (PRESS) curve (Analysis 3). No latent factor could be extracted to form a predictive model.

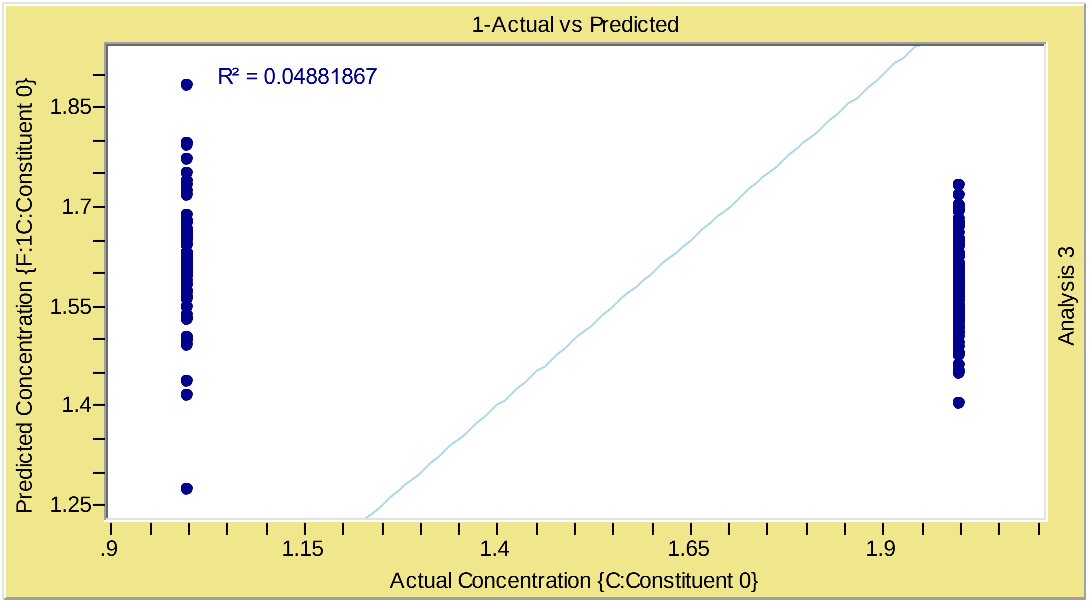


**Supplementary Figure 6** – Actual Vs Predicted plot. $R^2 = 0.049$. Constituent 1: uninfected; constituent 2: Sporozoite-infected (Analysis 3).
